# When Music Loses Its Pleasure: Hippocampal Cingulum White Matter as a Structural Mediator of Musical Reward Decline in Aging

**DOI:** 10.64898/2026.07.16.738989

**Authors:** Jinyu Wang, Nicholas Kathios, Benjamin Kubit, Kelsie Lopez, Ji Chul Kim, Edward Large, Stephanie Noble, Psyche Loui

## Abstract

Individuals’ ability to obtain pleasure from music, referred to as musical reward sensitivity, declines with age, yet the neural mechanisms underlying this decline remain unclear. In this study, we investigated musical reward sensitivity measured in 58 older adults and 131 young adults. Consistent with prior findings, young adults reported higher musical reward sensitivity than older adults (*p* < 0.05). To identify neuroanatomical predictors of musical reward sensitivity, we employed the elastic-net model to predict musical reward sensitivity using white matter microstructural properties and gray matter morphometric properties from the whole brain. In older adults, fractional anisotropy (FA) in the bilateral hippocampal cingulum (CGH) and external capsule (EC) reliably predicted individual differences in musical reward sensitivity. Moreover, FA in the right CGH significantly mediated the relationship between age and musical reward sensitivity in older adults. Notably, these associations were specific to musical reward sensitivity in older adults—that is, they did not replicate in young adults, and did not extend to general reward sensitivity. Together, these findings highlight the critical role of white matter integrity, particularly within the hippocampal-limbic pathways, in age-related changes in musical reward processing, and suggest a potential neurobiological target for interventions aimed at enhancing well-being in older adulthood.

**Key points:**

- Whole-brain machine learning identified the FA in the hippocampal cingulum and external capsule as robust predictors of musical reward sensitivity in older adults.
- FA in the right hippocampal cingulum mediated the association between age and musical reward sensitivity.
- These findings extend previous association-based studies by identifying hippocampal-limbic white matter as a key neural substrate supporting musical reward sensitivity decline during aging.

## Introduction

Music is one of the few rewards that people voluntarily seek throughout life, providing pleasure, emotional regulation, and social connection across cultures and across the lifespan (Dingle et al., 2021; Viola et al., 2023). However, recent evidence suggests that the ability to derive pleasure from music declines with age (Belfi et al., 2022), despite music remaining an important resource for promoting emotional well-being, reducing depression and stress, and supporting cognitive health in older adults (Böttcher et al., 2022; Creech et al., 2013; Xue et al., 2023). Why older adults experience less reward from music than others, and what neural mechanisms underlie this decline, remain largely unknown. Addressing this question is important not only for understanding how aging reshapes affective brain systems but also for identifying neurobiological targets that may help preserve emotional well-being in later life.

Neuroimaging studies have consistently shown that musical reward emerges through interactions between auditory, reward, and memory systems. Individuals with specific anhedonia, a condition where individuals do not experience pleasure from music despite normal hearing, musical perception, and normal pleasure from other rewards (Belfi & Loui, 2020), exhibit reduced activity of the nucleus accumbens and weaker functional connectivity between auditory cortex and ventral striatum during music listening (Martínez-Molina et al., 2016). Causal evidence further demonstrates that modulating frontostriatal circuitry using transcranial magnetic stimulation (TMS) alters the pleasure experienced during music listening (Mas-Herrero et al., 2018). Beyond classical auditory and reward regions, hippocampal activity has also been shown to encode musical uncertainty and surprise – computational features that strongly influence musical pleasure (Cheung et al., 2019). Together, these findings suggest that musical reward depends on distributed interaction among auditory, reward, and hippocampal networks.

Musical reward sensitivity has also been associated with variations in gray and white matter structure. For example, Hernández et al. (2019) reported negative associations between musical reward sensitivity and caudate and NAcc volumes. Diffusion Tensor Imaging (DTI) studies further indicate that white matter connectivity between the superior temporal gyrus (STG) and regions such as the insula and medial prefrontal cortex is positively associated with an individual’s experience of chills in response to music (Sachs et al., 2016). More broadly, white matter connectivity between frontal, striatal, and temporal regions (Martínez-Molina et al., 2019) is associated with BMRQ scores in the general population. Notably, a case study of musical anhedonia revealed decreased whiter matter volume but higher fractional anisotropy (FA) between auditory and striatal regions, particularly between left STG and left NAcc, suggesting complex microstructural alterations underlying impaired musical reward (Loui et al., 2017).

More recently, Matthews et al. (2024), using fixel-based analysis, reported positive associations between musical reward sensitivity and both fiber density and cross section in the right middle longitudinal fasciculus among musicians. Together, these studies suggest that structural brain organization contributes to individual differences in musical reward sensitivity. However, nearly all previous studies have relied on in-sample associations between brain measures and BMRQ scores. Although such approaches are valuable for generating detailed theories of mechanism, they offer limited ability to assess generalizability or predictive utility in new samples (Yarkoni & Westfall, 2017). Furthermore, most neuroimaging studies of musical reward sensitivity are based on predefined regions of interest (ROIs), with primary focus on auditory and reward-related regions. This strategy may overlook additional brain systems outside the auditory and reward network that contribute to musical reward processing. Consequently, it remains unclear whether structural brain measures can reliably predict musical reward sensitivity in unseen individuals or whether important brain systems outside canonical auditory-reward circuits also contribute.

Behavioral studies consistently show that musical reward sensitivity decreases with age (Belfi et al., 2022; Mannino et al., 2024; Mas-Herrero et al., 2013). For example, Belfi et al. (2022) examined individuals aged 20 to 85 years and reported a negative association between age and BMRQ scores. Parallel neuroimaging research has likewise demonstrated that aging alters neural processing of music. Compared with younger adults, older adults exhibit weaker correlation between activity in the temporal-mesolimbic reward network and subjective liking during music listening (Faber et al., 2023), greater ventral tegmental area (VTA) activation during music-evoked nostalgia (Hennessy et al., 2025), and reduced functional connectivity between auditory and reward networks both at rest and during music listening (2023). Together, these findings suggest that aging reshapes the functional brain networks supporting musical reward.

Meanwhile, age-related alterations also evident in brain structures, including reductions in gray matter volume and white matter integrity across frontal, temporal, limbic, and subcortical regions. For instance, compared to the young adults, older adults showed a widespread reduction in gray matter volume in the regions of the frontal, insular, cingulate cortices, and hippocampus, etc. (Farokhian et al., 2017; Nobis et al., 2019), as well as white matter volume reduction in several regions including thalamus, external capsule, internal capsule, cerebral peduncle, and the anterior cingulum (Farokhian et al., 2017; Giorgio et al., 2010; Pagani et al., 2008). Despite these paralell findings, an important question remains unanswered: do age-related structural brain changes help explain why musical reward sensitivity declines with age? To date, evidence addressing this question is remarkably limited, with only a case study examining white matter correlates of musical reward sensitivity in an older individual with musical anhedonia (Loui et al., 2017).

The present study aimed to determine whether age-related differences in brain structure contribute to individual differences in musical reward sensitivity. Specifically, we combined whole-brain structural neuroimaging with machine learning to identify neuroanatomical features that robustly predict musical reward sensitivity in unseen individuals and then examined whether these features statistically mediate the association between age and musical reward sensitivity.

Compared with previous ROI-based studies, our whole-brain approach incorporated gray matter morphometric measures from cortical and subcortical regions together with white matter microstructural properties, allowing the discovery of predictive brain regions beyond established auditory-reward circuits. By emphasizing out-of-sample prediction, the machine learning framework provides a more rigorous assessment of generalizable brain-behavior relationships than traditional correlational analyses (Gabrieli et al., 2015; Yarkoni & Westfall, 2017). Finally, because predictive models alone cannot determine how aging, brain structure, and behavior are related, mediation analyses were conducted to test whether the identified neuroanatomical features statistically explain age-related differences in musical reward sensitivity. Based on previous evidence implicating auditory, reward, and hippocampal systems in both musical reward and aging, we hypothesized that structural alterations within these networks would predict musical reward sensitivity and mediate the relationship between age and musical reward sensitivity. Ultimately, our goal was to explain why music loses its pleasure during aging by identifying structural brain pathways that link aging to changes in musical reward sensitivity.

## Materials & Methods

### Participants

A total of 58 older adults (age ≥ 50, age range: 54-89) and 131 young adults (age ≤ 25, age range: 18-25; see sample demographics are summarized in Table 1) were included in the present study. All participants provided MRI data, Barcelona Music Reward Questionnaire (BMRQ) scores, and Goldsmiths Musical Sophistication Index (Gold-MSI) scores in this study. These participants were recruited across multiple projects in the lab through a combination of online advertisements and community outreach events. Exclusion criteria were any history of psychotic or schizophrenic episodes, major neurologic diagnosis (Parkinson’s, stroke, brain injury, epilepsy), or other condition that might impair cognition or confound assessments, past or present substance abuse (alcohol and drugs), and common contraindications for MRI scanning. Participants filled out all these scales online through Qualtrics before MRI scan. All participants provided written informed consent prior to participation. The study was conducted in accordance with the Declaration of Helsinki and approved by the Institutional Review Board (IRB) of Northeastern University (IRB: 18-12-13 approved since May 2020 and IRB: 19-03-20 approved since March 2019).

**Table 1.** Sample characteristics.

| Variable | Older Adults (n = 58) | Young Adults (n = 131) |
| --- | --- | --- |
| Age (year), M(SD) | 66.84 (8.58) | 20.03 (1.80) |
| Birth Sex, n (%) |  |  |
| Female | 37 (63.79%) | 89 (67.94%) |
| Male | 21 (36.21%) | 40 (30.53%) |
| Unknown | 0 | 2 (1.53 %) |
| Gold-MSI Musical Training Score | 19.41 (9.57) | 23.34 (11.03) |

*Note.* M, mean; SD, standard deviation; Gold-MSI Musical Training Score, Musical Training subscale score from Goldsmiths Musical Sophistication Index.

### Behavioral Assessments

#### Barcelona Music Reward Questionnaire (BMRQ)

Individual differences in musical reward sensitivity are commonly measured using the Barcelona Music Reward Questionnaire (BMRQ), a 20-item self-report scale assessing five dimensions (subscales) of musical reward: Music Seeking (e.g., “I inform myself about music I like”), Emotion Evocation (e.g., “I like to listen to music that contains emotion”), Mood Regulation (e.g., “Music calms and relaxes me”), Sensory-Motor (e.g., “Music often makes me dance”), and Social Reward (e.g., “When I share music with someone I feel a special connection with that person”). Each item is rated on a 7-point Likert scale ranging from 1 (completely disagree) to 5 (completely agree). Item 2 and Item 5 are reversed items. The BMRQ has since been adapted and validated across multiple languages and cultures, supporting its reliability as a measure of musical reward sensitivity (Carraturo et al., 2025; Cetinbag-Kuzu et al., 2026; Honda et al., 2026; Lippolis et al., 2025; Mannino et al., 2024; Pereira & Bortoloti, 2025; Saliba et al., 2016; Wang et al., 2023). We used the English version of the BMRQ, which showed excellent reliability (Mas-Herrero et al., 2013).

#### Goldsmiths Musical Sophistication Index (Gold-MSI)

The Gold-MSI is a 38-item self-report tool that assesses musical behaviors, experiences, and skills across multiple dimensions (Müllensiefen et al., 2014). It comprises five subscales: Active Engagement (9 items, e.g., daily attentive music listening duration), Perceptual Abilities (9 items, e.g., identifying out-of-tune singing or playing), Musical Training (7 items, e.g., years of formal music theory training), Singing Abilities (7 items, e.g., accuracy in matching recorded notes while singing), and Emotions (6 items, e.g., selecting music for mood enhancement). Of those 38 items, 31 statements are rated on a 7-point Likert-type scale, and 7 questions use ordered response categories. We used the score of the musical training subscale as the index of musical training experience of participants. Each subscale score was computed by calculating the sum of item scores within the respective subscale.

#### Snaith-Hamilton Pleasure Scale (SHAPS)

The SHAPS is a 14-item scale measuring the degree to which a person experiences pleasure or anticipates pleasurable experiences. This scale covers four domains of general hedonic experience, including interest/pastimes, social interaction, sensory experience, and food/drink. Each of the items has a set of four response categories, including Definitely Agree (= 1), Agree (= 2), Disagree (= 3), and Definitely Disagree (= 4). A higher total score indicates higher levels of state anhedonia. The scale was constructed in such a way that cultural, gender, and age biases were kept to a minimum. The items of the scale relate to experiences likely to be encountered by most people (Snaith et al., 1995). 54 older participants and 14 young adults also responded to the Snaith-Hamilton Pleasure Scale (SHAPS) in the present study.

### MRI data acquisition and preprocessing

MRI data were collected using a 3T Siemens Prisma full-body scanner (Siemens Medical Solutions, Erlangen, Germany) with a 64-channel head coil at the Northeastern University Biomedical Imaging Center. For each participant, a rapid gradient echo (MPRAGE) T1-weighted structural image was acquired using a high-resolution 3D magnetization with the following parameters: field of view = 256 mm, 208 slices, resolution = 0.8 x 0.8 x 0.8 mm, time repetition (TR) = 2500 ms, time echos (TE) = 1.81, 3.6, 5.39, and 7.18 ms, flip angle = 8 degrees, phase encoding (PE) direction = anterior to posterior, IPAT mode = GRAPPA 2x, acquisition time = 8 min 22 sec. Whole-brain DWI data was acquired using a sing-shot spin-echo planar imaging sequence with the following parameters: field of view = 240 mm, 72 slices, resolution 2.00 mm isotropic, TR = 4770 ms, TE = 125 ms, flip angle = 78 degrees, PE direction = anterior to posterior, gradient directions = 64, b-values = 0, 1000, 2000 s/mm^2^, acquisition time = 10 min 34 sec.

### Gray-matter morphometric features

T1 images were processed and segmented using FreeSurfer 8.0.0, which segments bilateral subcortical structures and bilateral cortical regions in accordance with the Desikan-Killiany atlas (Desikan et al., 2006) and Aseg atlas (Fischl et al., 2002). Mean volume of subcortical brain structures and mean thickness of cortical regions were extracted. We finally included the cortical thickness (CT) of 68 regions and the subcortical volume (SV) from 28 subcortical structures as features (Detailed feature list is included in the supplementary materials).

### Fiber bundle features

We used the FSL 6.0.7 software to process DTI images, performing eddy current and head movement corrections, brain extraction, and tensor calculations to obtain fractional anisotropy (FA), axial diffusivity (AD), radial diffusivity (RD), and mean diffusivity (MD) maps. FA measures how directional the diffusion of water molecules is. AD measures the rate of water diffusion strictly parallel to the primary direction of the nerve fibers (axons). RD quantifies the rate of water diffusion perpendicular to the nerve fibers. MD measures the overall average rate at which water molecules diffuse through tissue (Le Bihan et al., 2001). From each participant’s DTI data, we extracted regional FA, AD, RD, and MD values using the JHU-ICBM-labels-1mm atlas, resulting in 50 features per participant for each type of metric (Detailed feature list is included in the supplementary materials).

### Construction of predictive models

To identify neuroanatomical predictors of musical reward sensitivity and general reward sensitivity, we built predictive models using a nested cross-validation framework based on Elastic Net regression (Zou & Hastie, 2005). Elastic Net regression model is a high-performance and stable machine learning model for neuroimaging data (Dobs et al., 2014; Jollans et al., 2019), which combines L1 norm regularization (Least Absolute Shrinkage and Selection Operator; LASSO; Tibshirani, 1996) and L2 norm regularization (Ridge; Hoerl & Kennard, 1970). The LASSO penalty produces sparse solutions by forcing the coefficients of less informative predictors to exactly zero, whereas the Ridge penalty stabilizes coefficient estimates in the presence of multicollinearity by shrinking correlated predictors toward similar values. Elastic Net therefore provides an effective compromise between feature selection and coefficient shrinkage, making it well suited for high-dimensional neuroimaging data with correlated predictors.

To reduce feature dimensionality and improve signal-to-noise ratio in the context of a modest sample size relative to the number of neuroanatomical predictors, we applied a correlation-based feature filtering step within each training fold prior to the Elastic Net regression following (Shen et al., 2017). In this step, features showing significant Pearson correlations with the target variable (BMRQ total score) at *p* < 0.05 were retained; if no features met this criterion, the top five features with the smallest *p*-values were included to ensure model stability. The preliminary feature filtering improved out-of-sample predictive performance in our dataset when compared with the pipeline without such preliminary feature filtering before the Elastic Net regression, and was therefore retained in the final pipeline.

We then implemented a nested Elastic Net regression (Varma & Simon, 2006) using a two-level cross-validation procedure to optimize hyperparameters and estimate generalization performance. In the inner loop, a 5-fold cross-validation was used to select the optimal regularization parameters (α and l₁ ratio) based on the coefficient of determination (R²). In the outer loop, a leave-one-out cross-validation (LOOCV) was conducted, where data from each participant served as the held-out test set once, and the model was trained on all remaining participants using the best hyperparameters determined from the inner loop. All preprocessing steps, including feature scaling, were fit exclusively within the training data of each fold to prevent information leakage.

Model performance was quantified using the coefficient of determination (R²), root mean squared error (RMSE), and mean absolute error (MAE) computed across all outer folds. To assess the statistical significance of model performance, we performed permutation testing (1000 iterations) by randomly shuffling BMRQ scores across participants and recomputing the nested LOOCV, thereby generating a null distribution for each performance metric. Empirical p-values were calculated as the proportion of permuted models that performed as well as or better than the true model. See Figure 1 for details.

**Figure 1.**
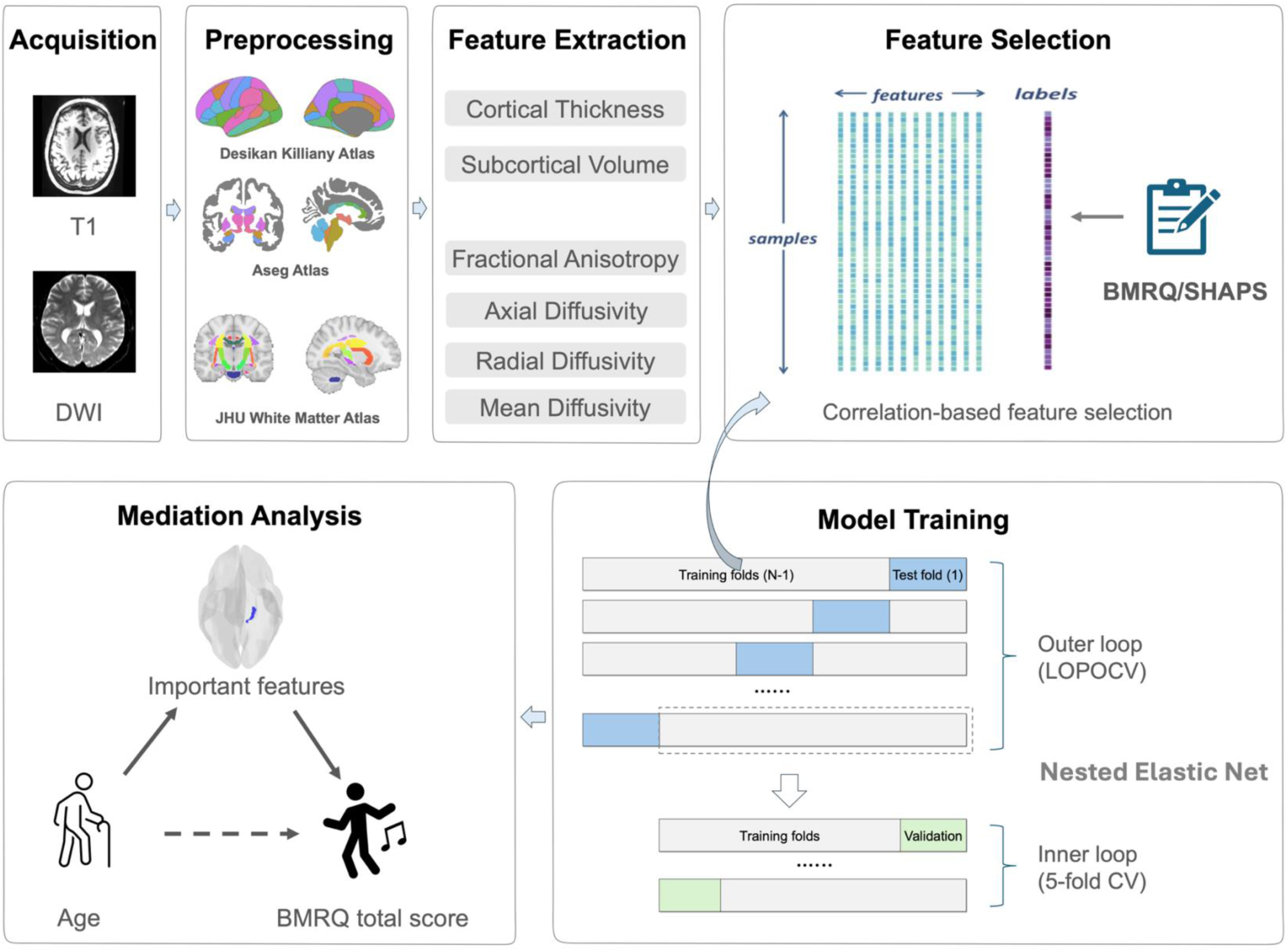
Schematic illustration of the overall workflow.

### Selection of important brain regions/tracts and mediation analysis

Feature importance was evaluated by extracting non-zero coefficients from each outer cross-validation fold of the nested elastic-net model. The frequency with which a given brain region or tract was selected as a non-zero predictor across folds was used to identify the most consistently contributing neuroanatomical features. To further investigate the potential mechanistic role of these features, mediation analyses were conducted using R (version 4.5.2). Specifically, we examined whether gray matter morphometric measures and white matter microstructural indices of identified regions/tracts mediated the association between age and musical reward sensitivity. A simple mediation model was specified, with age entered as the predictor (X) and BMRQ score as the outcome (Y). Bootstrap resampling procedures were leveraged to derive robust, bias-corrected, accelerated confidence intervals for the parameters of interest.

## Results

### Behavioral results

The assessment results are summarized in Table 2. For the older adults group, the mean BMRQ score and mean SHAPS score were 73.97 and 22.44, respectively. For the young adults group, the mean BMRQ score and mean SHAPS score were 79.68 and 22.86, respectively. The distributions of assessment scores are shown in Figure 2.

**Figure 2.**
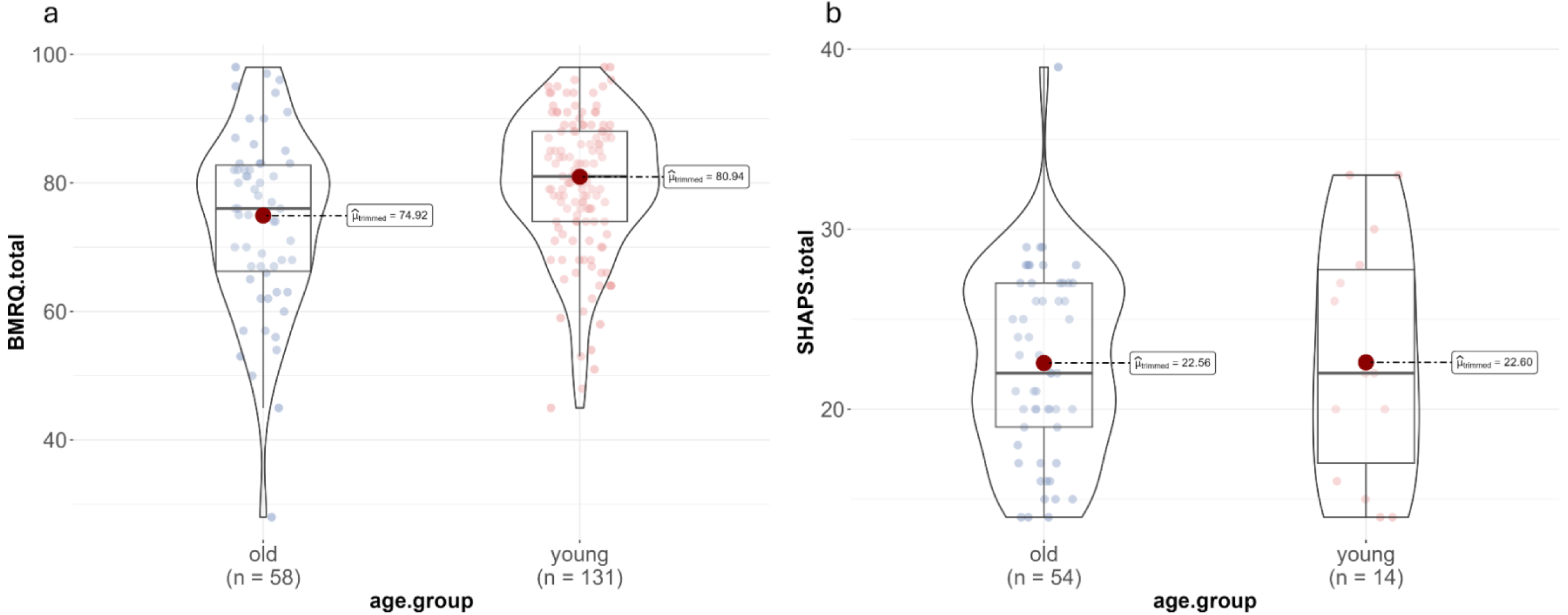
Distributions of musical reward sensitivity and general sensitivity in older and younger adults: (a) BMRQ score and (b) SHAPS score. *Note.* BMRQ.total, the total score of Barcelona Music Reward Questionnaire; SHAPS.total, the total score of Snaith-Hamilton Pleasure Scale; ^*μ*_trimmed_, the 20% trimmed mean used as a robust estimate of central tendency

**Table 2.** The assessment results of two scales.

| Group | BMRQ (M±SD) | SHAPS (M±SD) |
| --- | --- | --- |
| Older adults | 73.97±13.87 (n = 58) | 22.44±5.22 (n = 54) |
| Young adults | 79.70±10.88 (n = 131) | 22.86±6.76 (n = 14) |

*Note.* BMRQ, Barcelona Music Reward Questionnaire; SHAPS, Snaith-Hamilton Pleasure Scale; M, mean; SD, standard deviation.

#### BMRQ/SHAPS difference between the young and older adult groups

As the sample sizes of the young adults group and the older adults group are unbalanced, a bootstrap-based multiple regression (performed 5000 bootstrap resamples) was conducted to examine age-group differences in BMRQ scores, controlling for sex and musical training. The age-group coefficient (younger adults > older adults) was 3.59, with a bootstrap standard error of 1.89. A 95% percentile bootstrap CI ranged from -0.24 to 7.35, indicating a borderline effect. However, a two-tailed bootstrap significance test showed that only 3.30% of bootstrap replicates reversed the sign of the age effect (*p* < .05), suggesting that the age effect is statistically reliable: young adults reported higher BMRQ scores than older adults. The same bootstrap-based multiple regression (performed 5000 bootstrap resamples) was conducted to test whether there are age-group differences in the SHAPS score after controlling for sex, as previous studies also revealed significant gender effects on general reward sensitivity (Sun et al., 2025; Yang et al., 2022). The age-group coefficient (young vs. old) was 0.68, with a bootstrap standard error of 1.96. A 95% percentile bootstrap CI ranged from -3.32 to 4.52, indicating a borderline effect. A two-tailed bootstrap significance test showed that 37.22% of bootstrap replicates reversed the sign of the age effect (*p* = .37), suggesting that the age effect on SHAPS score is not significant.

#### Correlation between BMRQ/SHAPS and age

We calculated the correlation between age and BMRQ/SHAPS score within each group. Age was significantly associated with BMRQ score in the young adult group (*r* = -.256, *p* < .01), whereas the correlations were not significant in the older adult group (*r* = -.08, *p* = .578). The correlation between age and SHAPS score was not significant in both the older adults (*r* = .09, *p* = .541) and the young adults group (*r* = -.14, *p* = .637) (see Figure S1).

### Prediction performance

For each group (young adults and older adults), 6 Elastic-Net regression models were constructed based on 4 microstructural properties of white matter (FA, AD, RD, and MD) and 2 gray-matter morphometric properties (CT and SV), respectively. Individuals’ BMRQ scores were significantly predicted by the FA model in the older adults group, with predictive performance exceeding the level expected by chance (RMSE = 11.99; MAE = 9.77; *r* = .40, permutation test *p* < .05, FDR corrected over 6 models) (see Figure 3). However, the FA model did not perform significantly better than chance level in the young adults group (RMSE = 10.62; MAE = 8.44; *r* = .16, *p* = .144, FDR corrected over 6 models). Models constructed based on other properties (AD, RD, MD, CT, and SV) did not achieve predictive accuracies significantly greater than chance in permutation tests. The prediction results of all models are presented in Table 3a. The same models were also constructed to predict individuals’ SHAPS scores. However, the predictive performance of all 6 models did not perform significantly better than the level expected by chance, either in the older adults or the young adults. All models’ performance metrics are summarized in Table 3b.

**Figure 3.**
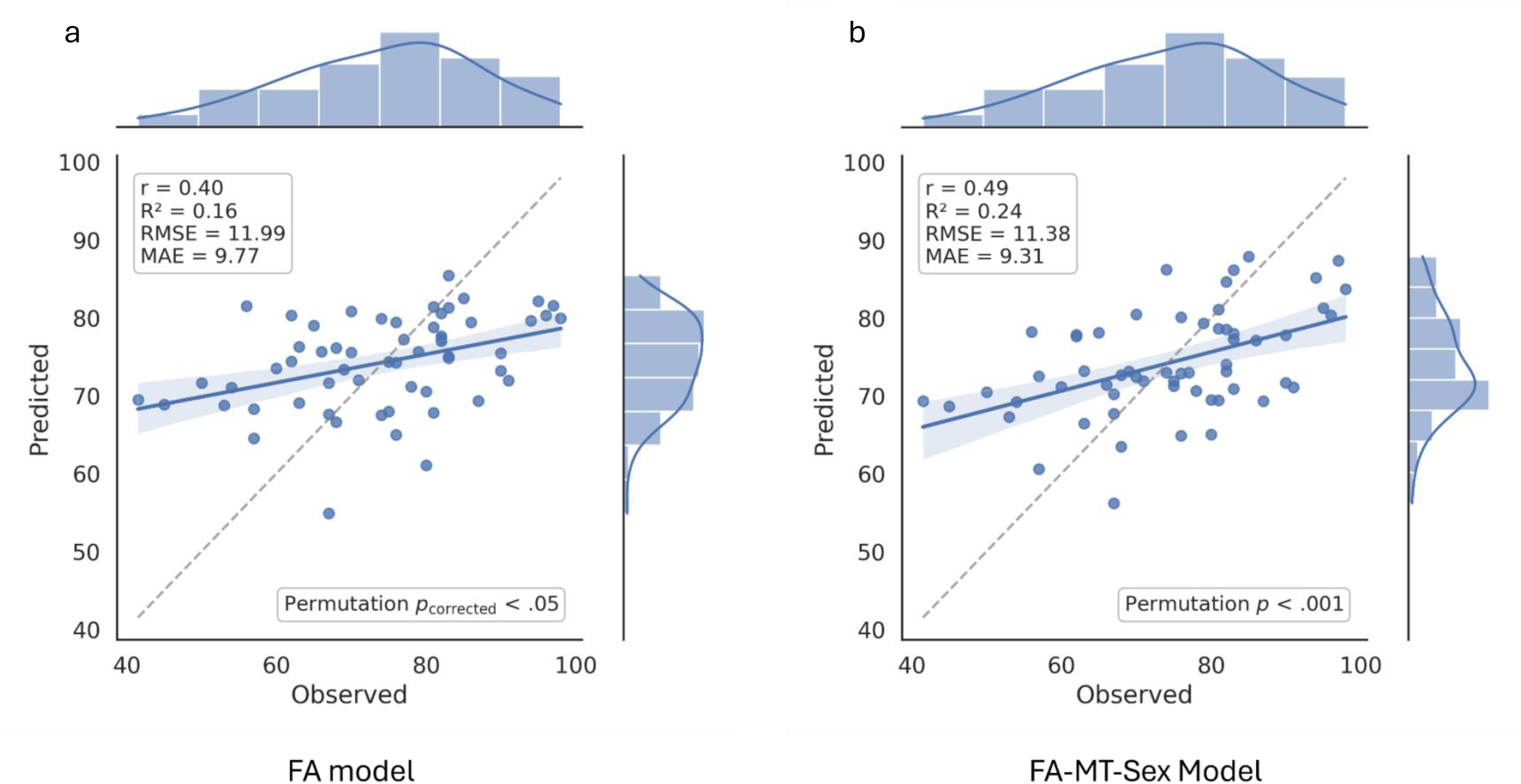
The relationship between observed and predicted BMRQ score in the FA model (a) and the FA-MT-Sex model (b) in older adults. *Note.* FA model = elastic-net model using FA features only; FA-MT-Sex model = elastic-net model using FA features together with musical training score and sex as predictors. Each point represents one participant. The solid blue line indicates the least-squares regression fit between observed and predicted BMRQ scores, and the shade area represents the 95% confidence interval of the regression line. The dashed gray diagonal line represents perfect agreement between observed and predicted scores. Histograms and density curves along the top and right margins show the distribution of observed and predicted BMRQ scores, respectively. *r*, Pearson’s correlation coefficient; R², coefficient of determination; RMSE, root mean squared error; MAE, mean absolute error; Permutation *p_corrected_*, *p* value of permutation tests after False Discovery Rate (FDR) correction; Permutation *p*, *p* value of permutation tests.

**Table 3a.**
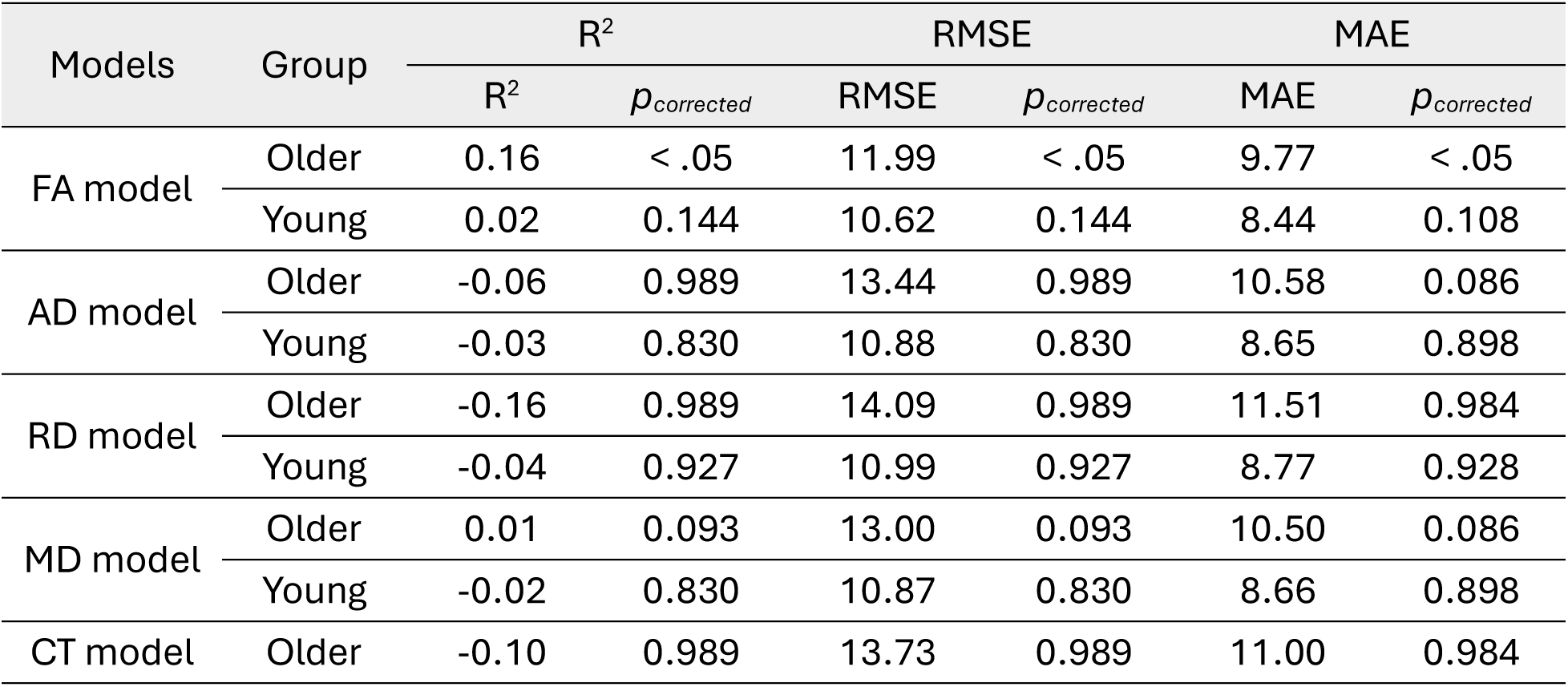
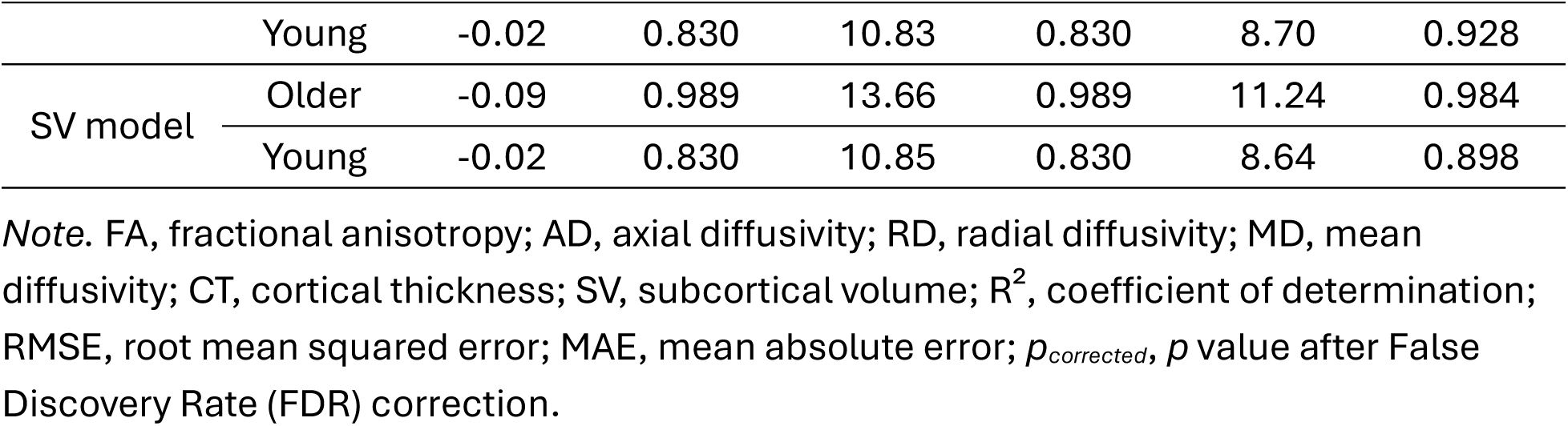
Prediction results of 6 Elastic-Net regression models predicting BMRQ score for young adults and older adults.

**Table 3b.**
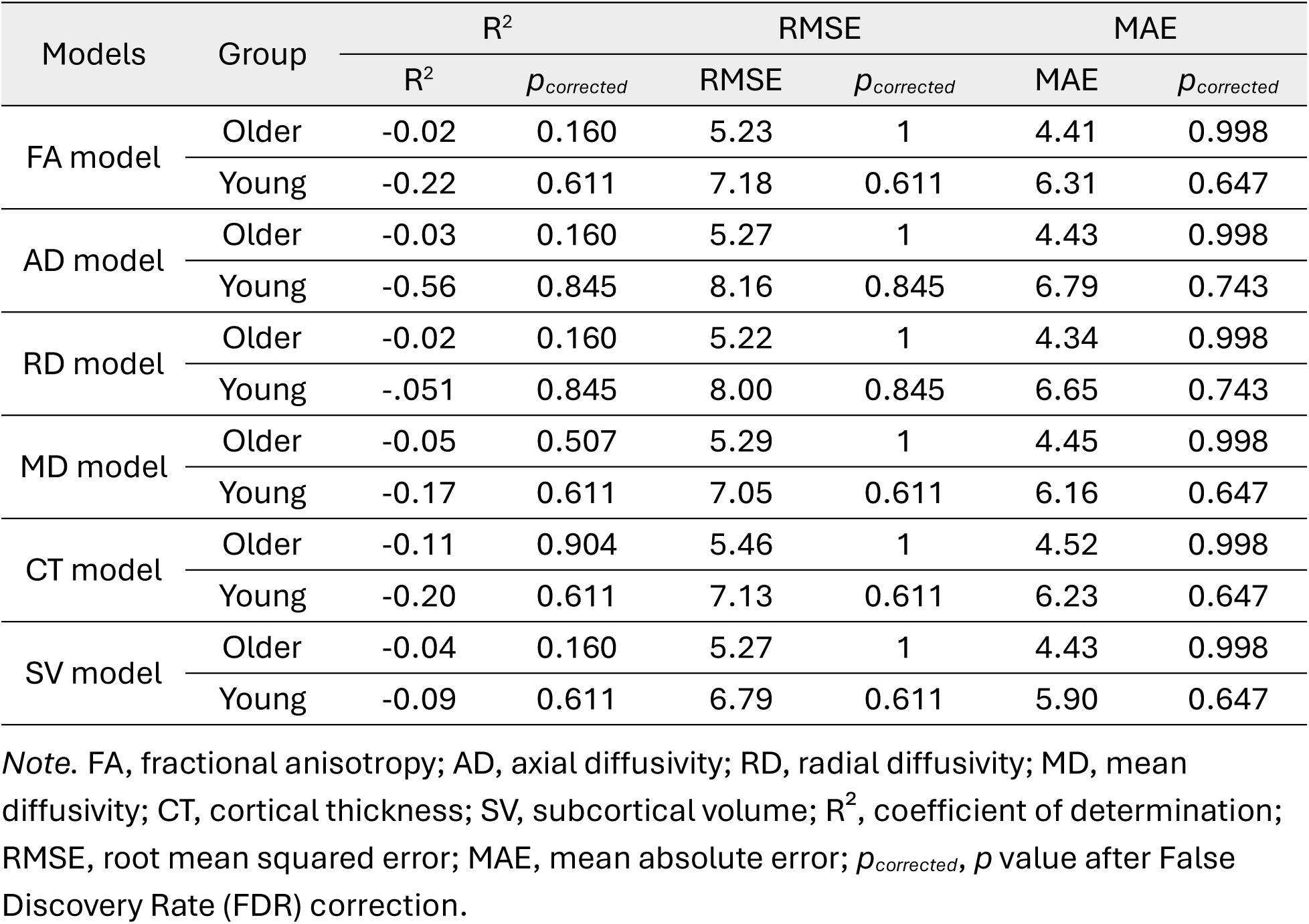
Prediction results of 6 Elastic-Net regression models predicting SHAPS score for young adults and older adults.

Differences in musical reward sensitivity have been linked to both musical training (Carraturo et al., 2025; Honda et al., 2023; Mas-Herrero et al., 2013) and sex (Carraturo et al., 2025; Honda et al., 2023; Mas-Herrero et al., 2013; Wang et al., 2023). As such, we reconstructed the FA model with additional covariates for sex and musical training (as measured by Gold-MSI musical training scale). To evaluate the additive predictive value of FA features, we compared a baseline model including only covariates (sex and musical training score) with a full model (FA-MT-Sex model) including FA features. The full model significantly predicted individuals’ BMRQ scores (RMSE = 11.38; MAE = 9.31; *r* = .49, *p* < .001) and demonstrated improved predictive performance relative to the baseline model (RMSE = 12.66; MAE = 10.09; *r* = .26, *p* < .01) (see Figure S2). This indicates that FA features provide additional explanatory power beyond sex and musical training.

### Tracts predictive of musical reward sensitivity

For the FA model in older adults, 4 predictors survived in all 58 LOOCV loops, including the left and right hippocampal cingulum (CGH) tracts (visualized in Figure 4a), the left external capsule (EC) tract (Figure 4b), and the left superior longitudinal fasciculus (SLF) tract (Figure S3b). The frequencies of tracts that appeared as non-zero predictors among all 58 older adults in the FA model are listed in Table 4a. When including musical training score and sex in the model (FA-MT-Sex model), the right hippocampal cingulum tract, the left hippocampal cingulum tract, and the left external capsule tract still survived in all 58 LOOCV loops, whereas the left SLF no longer appeared consistently as a non-zero predictor across all LOOCV loops. This finding suggests that the predictive contribution of the left SLF may be partially accounted for by individual differences in musical training and/or sex, whereas the bilateral CGH and left EC remained robust predictors independent of these covariates. In addition, musical training score appeared as a non-zero predictor among all 58 older adults. The frequencies of tracts appeared as non-zero predictors among all 58 older adults in the FA-MT-Sex model are listed in Table 4b. The FA values of these tracts and musical training score were all correlated to the BMRQ total score (*p*s < .01) (See Figure 5).

**Figure 4.**
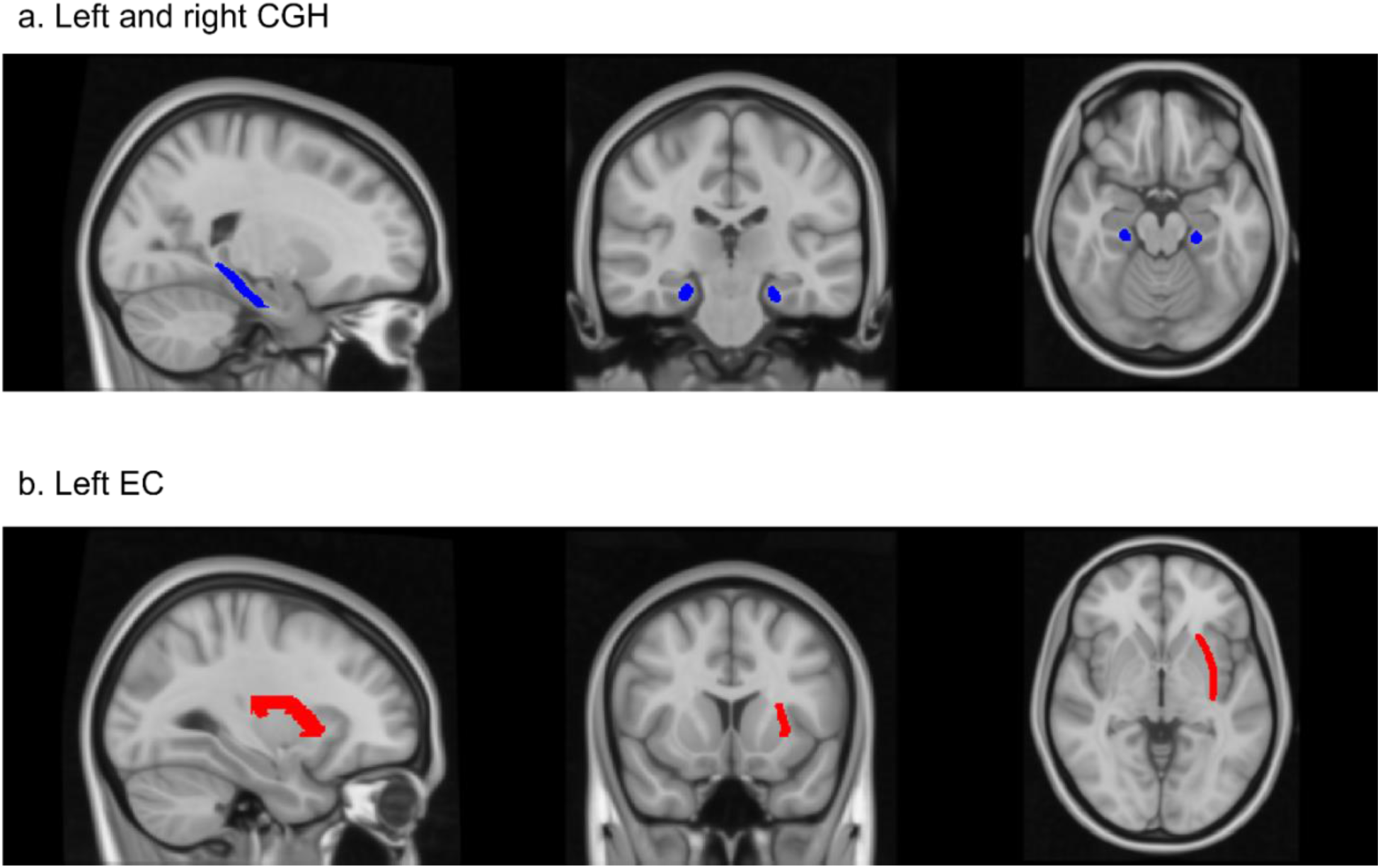
Fiber tracts predictive of musical reward sensitivity for the FA-MT-Sex model. *Note*. a, the left and right hippocampal cingulum tract; b, the left external capsule tract; c, the left superior longitudinal fasciculus tract.

**Figure 5.**
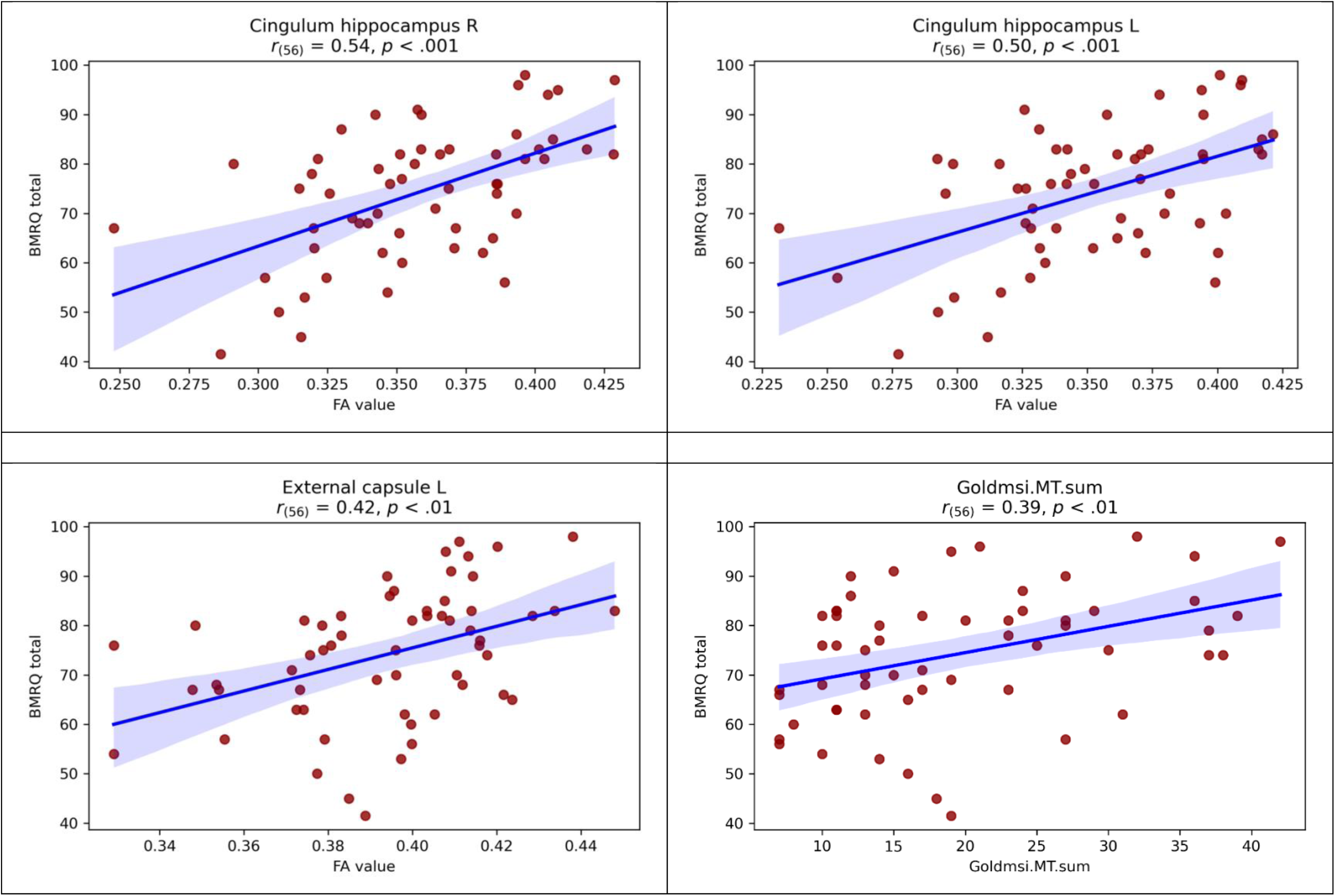
Correlations between FAs of important tracts/musical training score and BMRQ score. *Note.* Cingulum hippocampus R, the right hippocampal cingulum tract; Cingulum hippocampus L, the left hippocampal cingulum tract; External capsule L, the left external capsule; Goldmsi.MT.sum, musical training score from Gold-MSI; Superior longitudinal fasciculus L, the left superior longitudinal fasciculus; BMRQ total, the total score of Barcelona Music Reward Questionnaire.

**Table 4a.** The frequencies of tracts appeared as non-zero predictors among all 58 older adults in FA model predicting BMRQ.

| Tract | Frequency | *Pearson correlation between FA and BMRQ* |  |
| --- | --- | --- | --- |
|  |  | *r* | *p* |
| Cingulum_hippocampus_R | 58 | 0.54 | < .001 |
| Cingulum_hippocampus_L | 58 | 0.50 | < .001 |
| External_capsule_L | 58 | 0.42 | < .01 |
| Superior_longitudinal_fasciculus_L | 58 | 0.31 | < .05 |
| Cerebral_peduncle_L | 31 | 0.37 | < .01 |
| External_capsule_R | 30 | 0.36 | < .01 |
| Corticospinal_tract_L | 29 | 0.27 | < .05 |
| Sagittal_stratum_R | 26 | 0.27 | < .05 |
| Fornix_cres_Stria_terminalis_L | 26 | 0.35 | < .01 |
| Fornix_cres_Stria_terminalis_R | 24 | 0.33 | < .05 |
| Retrolenticular_part_of_internal_capsule_L | 21 | 0.35 | < .01 |
| Superior_longitudinal_fasciculus_R | 21 | 0.27 | < .05 |
| Cerebral_peduncle_R | 16 | 0.29 | < .05 |
| Retrolenticular_part_of_internal_capsule_R | 8 | 0.31 | < .05 |
| Posterior_thalamic_radiation_R | 5 | 0.25 | .054 |
| Middle_cerebellar_peduncle | 5 | 0.24 | .065 |
| Posterior_thalamic_radiation_L | 2 | 0.25 | .056 |
| Inferior_fronto_occipital_fasciculus_L | 1 | 0.21 | .105 |
| Posterior_corona_radiata_L | 1 | 0.16 | .221 |

Note. FA, fractional anisotropy; BMRQ, Barcelona Music Reward Questionnaire. \_R, right hemisphere, \_L, left hemisphere.

**Table 4b.**
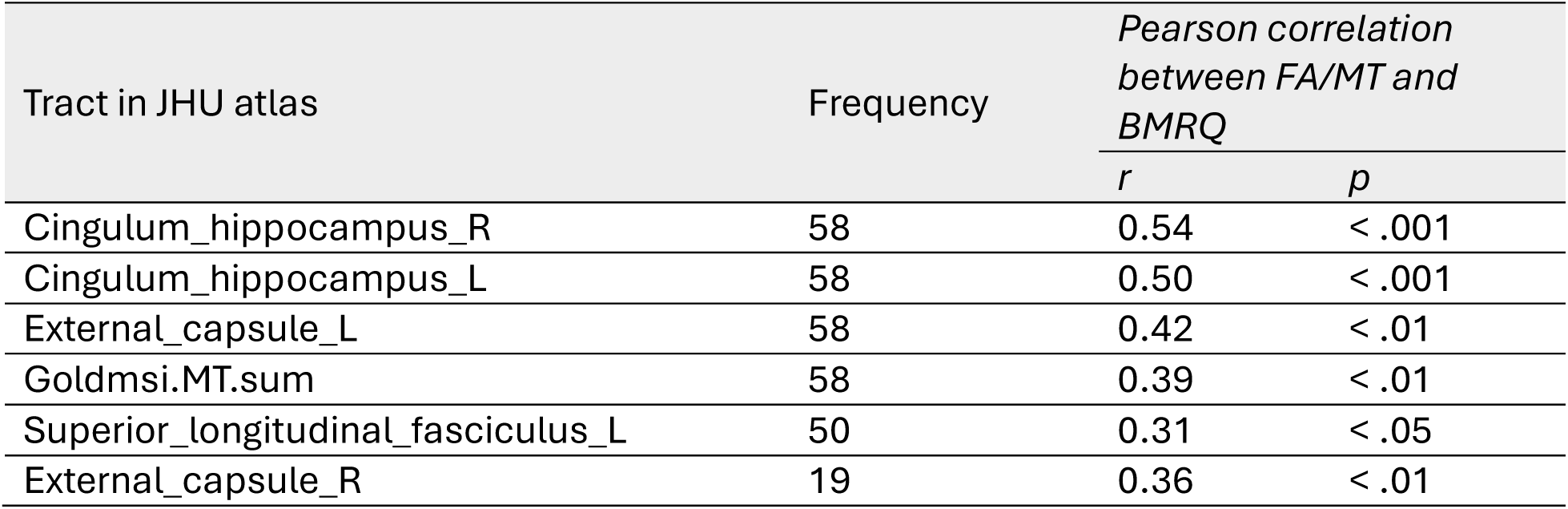
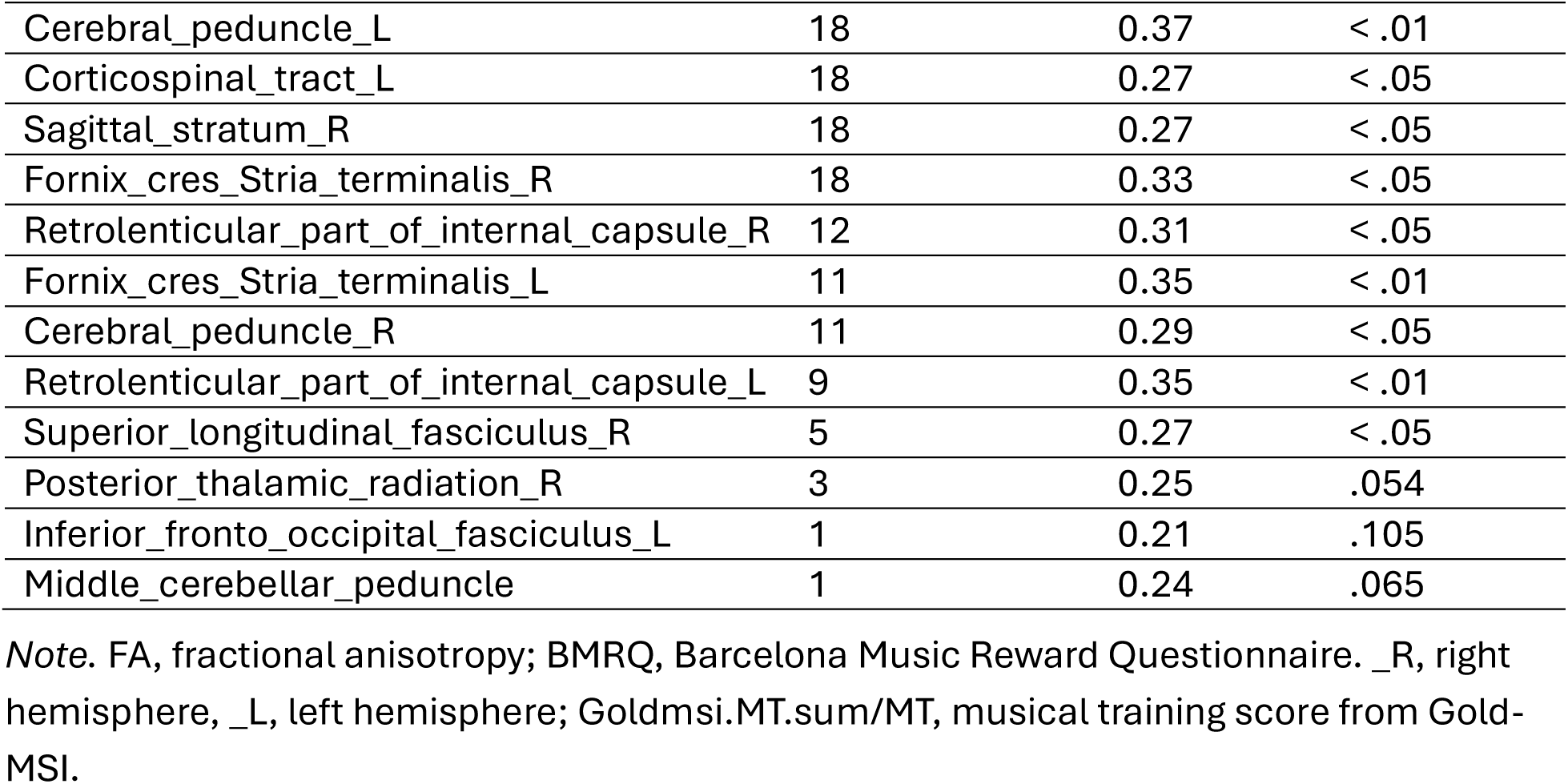
The frequencies of tracts appeared as non-zero predictors among all 58 older adults in FA-MT-Sex model predicting BMRQ.

| Tract in JHU atlas | Frequency | *Pearson correlation between FA/MT and BMRQ* |  |
| --- | --- | --- | --- |
|  |  | *r* | *p* |
| Cingulum_hippocampus_R | 58 | 0.54 | < .001 |
| Cingulum_hippocampus_L | 58 | 0.50 | < .001 |
| External_capsule_L | 58 | 0.42 | < .01 |
| Goldmsi.MT.sum | 58 | 0.39 | < .01 |
| Superior_longitudinal_fasciculus_L | 50 | 0.31 | < .05 |
| External_capsule_R | 19 | 0.36 | < .01 |
| Cerebral_peduncle_L | 18 | 0.37 | < .01 |
| Corticospinal_tract_L | 18 | 0.27 | < .05 |
| Sagittal_stratum_R | 18 | 0.27 | < .05 |
| Fornix_cres_Stria_terminalis_R | 18 | 0.33 | < .05 |
| Retrolenticular_part_of_internal_capsule_R | 12 | 0.31 | < .05 |
| Fornix_cres_Stria_terminalis_L | 11 | 0.35 | < .01 |
| Cerebral_peduncle_R | 11 | 0.29 | < .05 |
| Retrolenticular_part_of_internal_capsule_L | 9 | 0.35 | < .01 |
| Superior_longitudinal_fasciculus_R | 5 | 0.27 | < .05 |
| Posterior_thalamic_radiation_R | 3 | 0.25 | .054 |
| Inferior_fronto_occipital_fasciculus_L | 1 | 0.21 | .105 |
| Middle_cerebellar_peduncle | 1 | 0.24 | .065 |

*Note.* FA, fractional anisotropy; BMRQ, Barcelona Music Reward Questionnaire. \_R, right hemisphere, \_L, left hemisphere; Goldmsi.MT.sum/MT, musical training score from Gold-MSI.

### FA in the hippocampal cingulum tract mediates the relationship between age and musical reward sensitivity in older adults

A mediation model was developed to explore the relationship among age, important features (including FA in the right hippocampal cingulum tract, the left hippocampal cingulum tract, and the left external capsule tract), and musical reward sensitivity. We hypothesized that these features would mediate the relationship between age and musical reward sensitivity in older adults. To test this hypothesis, we constructed a structural equation model (SEM). The model included age as a predictor and the BMRQ score as a dependent variable. Table 5 and Figure 6 display the statistics of mediation effects in the SEM. The SEM revealed a significant indirect effect of age on musical reward sensitivity through FA in the right hippocampal cingulum tract (standardized beta = -0.24, *p* = 0.010). Older age was associated with reduced FA in this tract (standardized beta = -0.39, *p* = 0.002), and lower FA was in turn associated with reduced musical reward sensitivity (standardized beta = 0.61, *p* < .001). The direct effect of age on BMRQ was not significant after accounting for FA, indicating that age is associated with variation in BMRQ through age-related differences in FA of the right hippocampal cingulum, even though age itself does not show a significant overall association with BMRQ.

**Figure 6.**
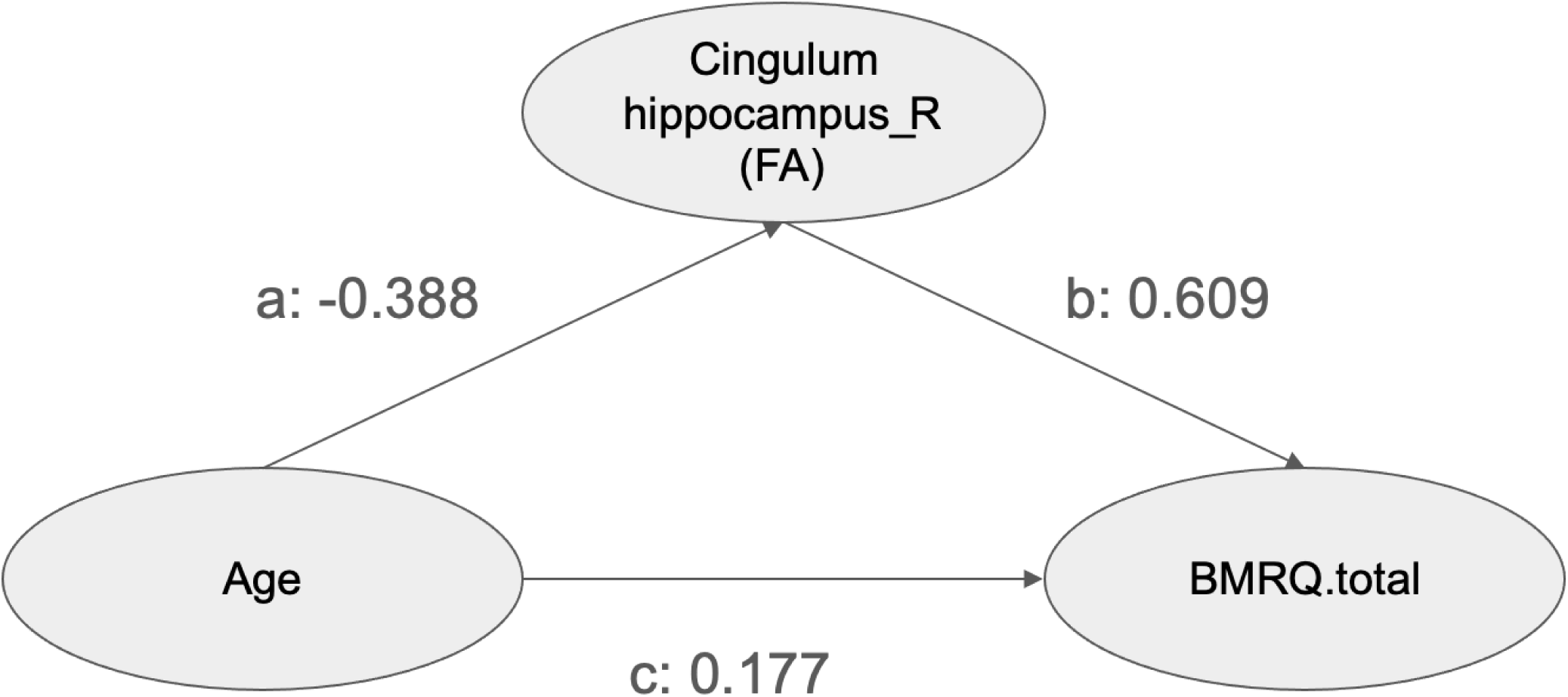
Path diagram for the mediation analysis using structural equation modeling (SEM). *Note.* Standardized coefficients are shown for each path. The bootstrap statistical significance of the indirect path is presented in Table 3. Results of the proposed model confirm that the age is negatively associated with BMRQ total score, through lower FA value in the right hippocampal cingulum tract.

**Table 5.**
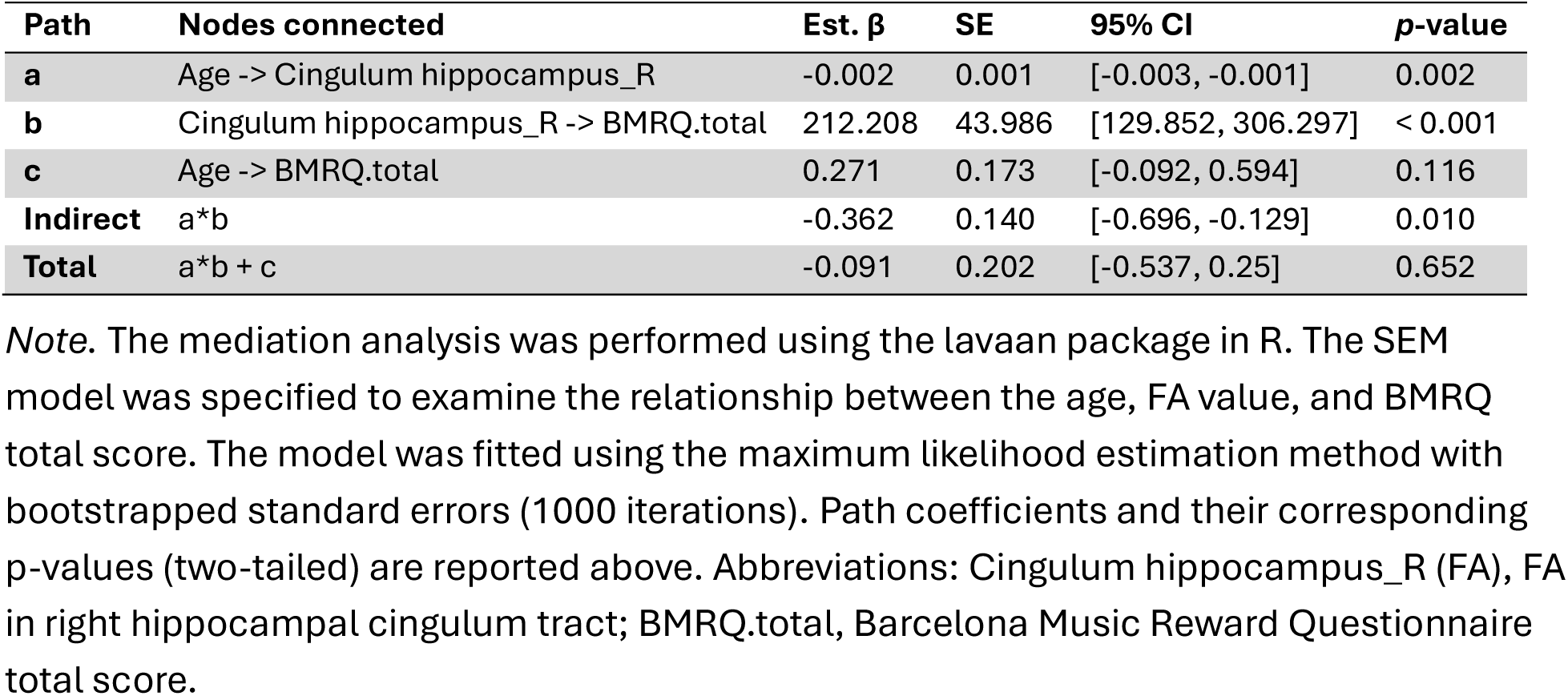
Results from the structural equation model (SEM)

## Discussion

The present study employed machine learning to predict musical reward sensitivity from white matter microstructural properties and gray matter morphometric properties in both older and young adults. In older adults, inter-individual variability in the white matter integrity within the right hippocampal cingulum tract, the left hippocampal cingulum tract, and the left external capsule significantly contributed to individual differences in musical reward sensitivity in older adults. Specifically, the FA values of these tracts were positively associated with musical reward sensitivity. Furthermore, FA in the right hippocampal cingulum tract mediated the association between age and musical reward sensitivity in older adults, suggesting a potential mechanistic pathway linking age-related white matter decline to reduced musical reward responsiveness.

Notably, these associations were not observed in young adults and did not generalize to general reward sensitivity as measured by the SHAPS. Together, these findings highlight a selective contribution of the hippocampal cingulum and external capsule to musical reward sensitivity in aging, underscoring the importance of white matter integrity within memory–emotion integration pathways in supporting music-related reward processes later in life.

We found that the FA value of the external capsule (EC) is an important feature contributing to musical reward sensitivity in older adults. FA is a sensitive diffusion MRI metric that reflects the degree of directionally constrained water diffusion within white matter, providing an index of white matter integrity (Alexander et al., 2007; Basser & Pierpaoli, 2011; Madden et al., 2012).

Higher FA values are associated with greater fiber coherence, higher axonal density, and intact myelination, whereas lower FA is observed in conditions with microstructural disruption (such as demyelination, axonal injury, or disorganized fiber architecture) (Sammer et al., 2022; Zhang et al., 2017). Specifically, the FA value of EC is positively associated with musical reward sensitivity. EC, located lateral to the internal capsule, is a thin layer of white matter fiber in the brain that traverses between the putamen and claustrum. It serves as a conduit for association fibers, including pathways associated with the inferior fronto-occipital fasciculus (IFOF), and the uncinate fasciculus (UF), which are crucial for communication between different cortical regions, particularly in the ventral pathway, linking frontal, temporal, and parietal lobes (Biswas et al., 2022; Mori et al., 2008). This connectivity supports multiple functions including language processing, emotion, and decision making (Dick et al., 2014; Von Der Heide et al., 2013).

The inclusion of EC as a key feature in our model is consistent with previous findings that emphasized the insula and its connected structures in music reward. The anterior insula (AIns) is repeatedly implicated in music emotion and reward processing (Belfi & Loui, 2020). For instance, larger tract volume between STG and insula was shown in individuals who frequently experience highly intense emotional responses to music compared to matched controls (Sachs et al., 2016). It is also a significant predictor of BMRQ score from a prior case study (Loui et al., 2017). The EC neighbors the insula and claustrum and contains fiber pathways connecting cortical areas to subcortical nuclei. For example, anatomical studies have shown that projections from the temporal lobe (including limbic regions like the entorhinal and perirhinal cortices) reach the nucleus accumbens (ventral striatum) via the external capsule (Groenewegen et al., 1982). In other words, the external capsule can carry signals from auditory and limbic related cortices to the NAcc, a key structure in reward processing. Previous findings on musical reward sensitivity and white matter tracts converged on the connectivity between auditory regions and the brain’s reward circuitry (insula, nucleus accumbens, orbitofrontal/mPFC) (Loui et al., 2017; Martínez-Molina et al., 2019; Matthews et al., 2024; Sachs et al., 2016). Our finding of the left EC’s importance is therefore quite consistent with the known auditory-reward circuit – it likely reflects the structural pathway through which information from frontal/insular and temporal regions accesses the striatal reward system. This complements earlier DTI results, showing that STG-insula-NAcc connectivity relates to musical reward (Loui et al., 2017), as the external capsule is one route for connecting insula/claustrum regions with the NAcc and other basal forebrain structures. In summary, a stronger or more efficiently connected left external capsule may facilitate the integration of auditory-emotional signals with dopaminergic reward processing.

In addition, we also found that the FA values of the right and left hippocampal cingulum tracts are important features contributing to individuals’ musical reward sensitivity in older adults. Specifically, the FA values of the right and left hippocampal cingulum tracts are positively correlated with the BMRQ score. These results indicated that musical reward sensitivity is not only linked to the auditory-reward circuit but also affected by the integrity of white matter pathways connecting the hippocampus and the cingulate gyrus in older adults. Although our sample size for young adults was larger, these relationships between FA and BMRQ were not detected in young adults. One possible explanation is the narrower distribution of FA values observed in young adults relative to older adults (see Figure S4). A wider distribution of predictor values may facilitate the detection of brain–behavior relationships, whereas restricted variability can attenuate predictive performance. This pattern is consistent with previous studies suggesting that white matter aging is characterized by substantial inter-individual variability and heterogeneous tract-specific age-related changes, resulting in greater heterogeneity of white matter microstructure in older adults than in younger adults (Matijevic & Ryan, 2025; Mendez Colmenares et al., 2023).

The hippocampal cingulum (CGH) tract is the hippocampal portion of the cingulum bundle, located inferior to the axial level of the splenium of the corpus callosum. It is part of the brain’s limbic system, connecting the hippocampus to the cingulate gyrus (Mori et al., 2008; Schmahmann & Pandya, 2009). The CGH facilitates communication for memory, emotion, and executive functions. Previous studies indicated that atrophy or microstructural changes (like reduced FA) in the CGH are linked to memory decline in aging and conditions like Alzheimer’s disease. For instance, the RD in CGH was found to be a significant independent predictor of episodic memory performance in older adults (Alm et al., 2022). Furthermore, focal changes in white matter tract properties in the right CGH also predicted memory and cognitive functioning in individuals with amnestic mild cognitive impairment (aMCI) (Gozdas et al., 2020).

Alterations in the CGH are also implicated in bipolar disorder. Kafantaris et al. (2017) found that patients with bipolar disorder demonstrated significantly lower FA compared to healthy individuals, and increased FA has been found as a biomarker of resilience to mood disorders (Roberts et al., 2022). Importantly, increased connectivity between the hippocampus and cingulate has been associated with focusing attention on music-evoked autobiographical memory (Kubit & Janata, 2018). Furthermore, Cheung et al. (2019) found that the interaction between uncertainty and surprise significantly modulated hippocampal activity during music listening, suggesting that the hippocampus contributes to both memory-related and predictive mechanisms underlying musical reward. Given the role that CGH plays in supporting memory and emotion, the association between white matter integrity in the hippocampal cingulum and musical reward sensitivity found in this study may reflect that individuals’ musical reward sensitivity is also supported by memory. Our finding that FA in the right hippocampal cingulum mediates the relationship between age and musical reward sensitivity in older adults helps explain why musical reward sensitivity changes in the aging process. As age increases the FA value in the hippocampal cingulum decreases, resulting in lower musical reward sensitivity.

Taken together, our results expand on previous structural connectivity studies to investigate the underlying mechanisms of individual differences in musical reward sensitivity. We identified the white matter tracts – the left EC and CGH, which are predictive of older adults’ musical reward sensitivity using whole-brain features and predictive models, and revealed the mediation effect of the FA value in the right CGH on the relationship between age and musical reward sensitivity in older adults.

The present study has several strengths. A major strength of this study is the focus on out-of-sample prediction with the elastic net. This approach enables more generalizable forecasting of musical reward sensitivity in new subjects from white-matter structures and provides an important extension to prior association-based studies. Our use of whole-brain analysis instead of predefined regions of interest on the auditory and reward regions enables us to combine information across multiple areas of the brain, simultaneously, consistent with an increasing understanding that brain mechanisms underlying behavior are multivariate and widespread across the brain (Noble et al., 2024), which provides us with a more comprehensive understanding of musical reward sensitivity. By including young adults’ data and general reward sensitivity data in our analysis, we can test if these findings are unique to aging and musical reward sensitivity specifically.

There are also limitations in our study. There are relatively large differences in the sample size between the older adults and the young adults group. The number of participants who have the musical reward sensitivity score and the general reward sensitivity score is also different. Such imbalances may result in unequal statistical power across groups, making it difficult to directly compare between groups and variables. Additionally, our study assessed the musical reward sensitivity using only a self-reported scale, which may be influenced by subjective interpretation and response bias and may not fully capture behavioral or physiological aspects of musical reward sensitivity. Future studies with more balanced sample sizes and multimodal assessments of musical reward sensitivity, including behavioral and physiological measures, are needed to improve our understanding of the construct of musical reward sensitivity and how it is instantiated in the brain.

## Conclusion

The current study investigated the neural underpinnings of musical reward sensitivity changes in the aging process using nested elastic-net regression models. Our findings indicate that white matter integrity in the bilateral CGH and the left EC predicts older adults’ musical reward sensitivity. Importantly, the FA value in the right CGH mediates the relationship between age and musical reward sensitivity in older adults. These findings expand the understanding of the neural mechanism of musical reward sensitivity and how the aging process affects individuals’ ability to perceive pleasure from music. More broadly, identifying white matter pathways associated with musical reward sensitivity may help inform personalized music-based interventions by clarifying which neural systems support rewarding responses to music in aging populations.

## Acknowledgments

The authors would like to thank all the individuals who participated in this study.

## Data availability statement

The data and code used in the present study are available from the corresponding author upon request.

## Supplementary Material

**Figure S1.**
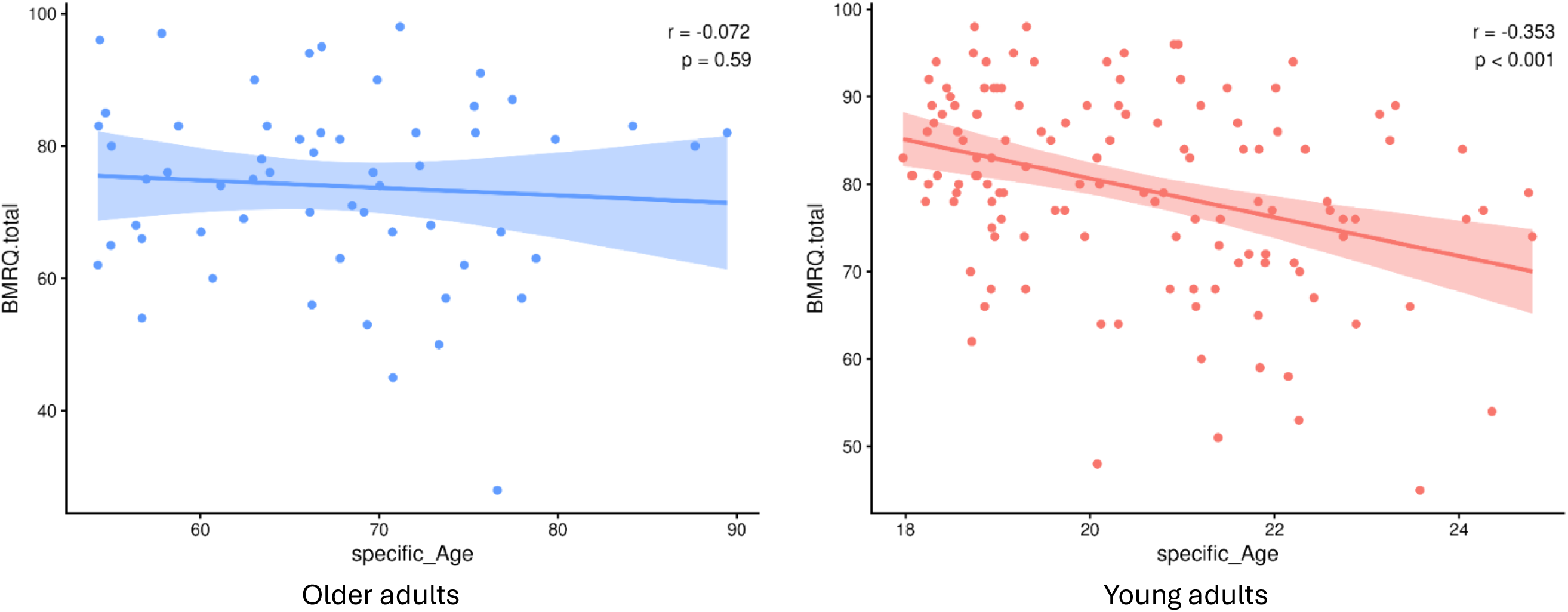
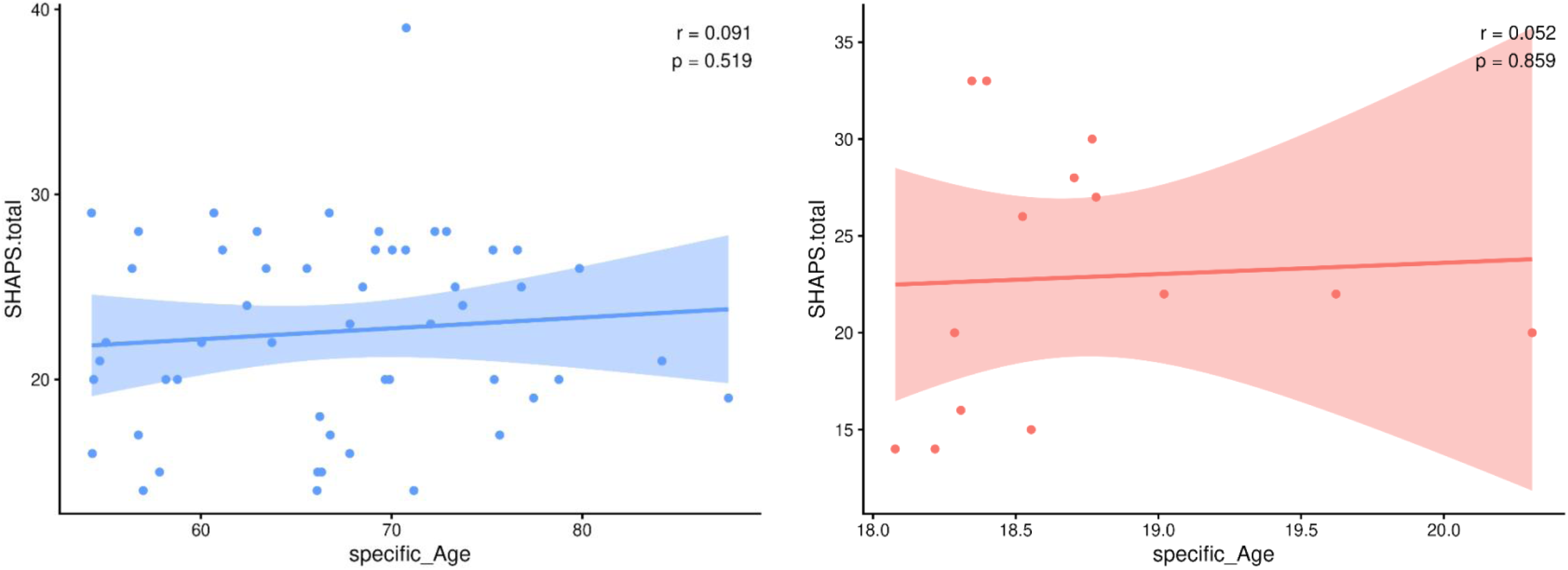
Correlations between age and BMRQ/SHAPS scores. *Note.* BMRQ.total, the total score of Barcelona Music Reward Questionnaire; SHAPS.total, the total score of Snaith-Hamilton Pleasure Scale; *r*, correlation coefficient.

**Figure S2.**
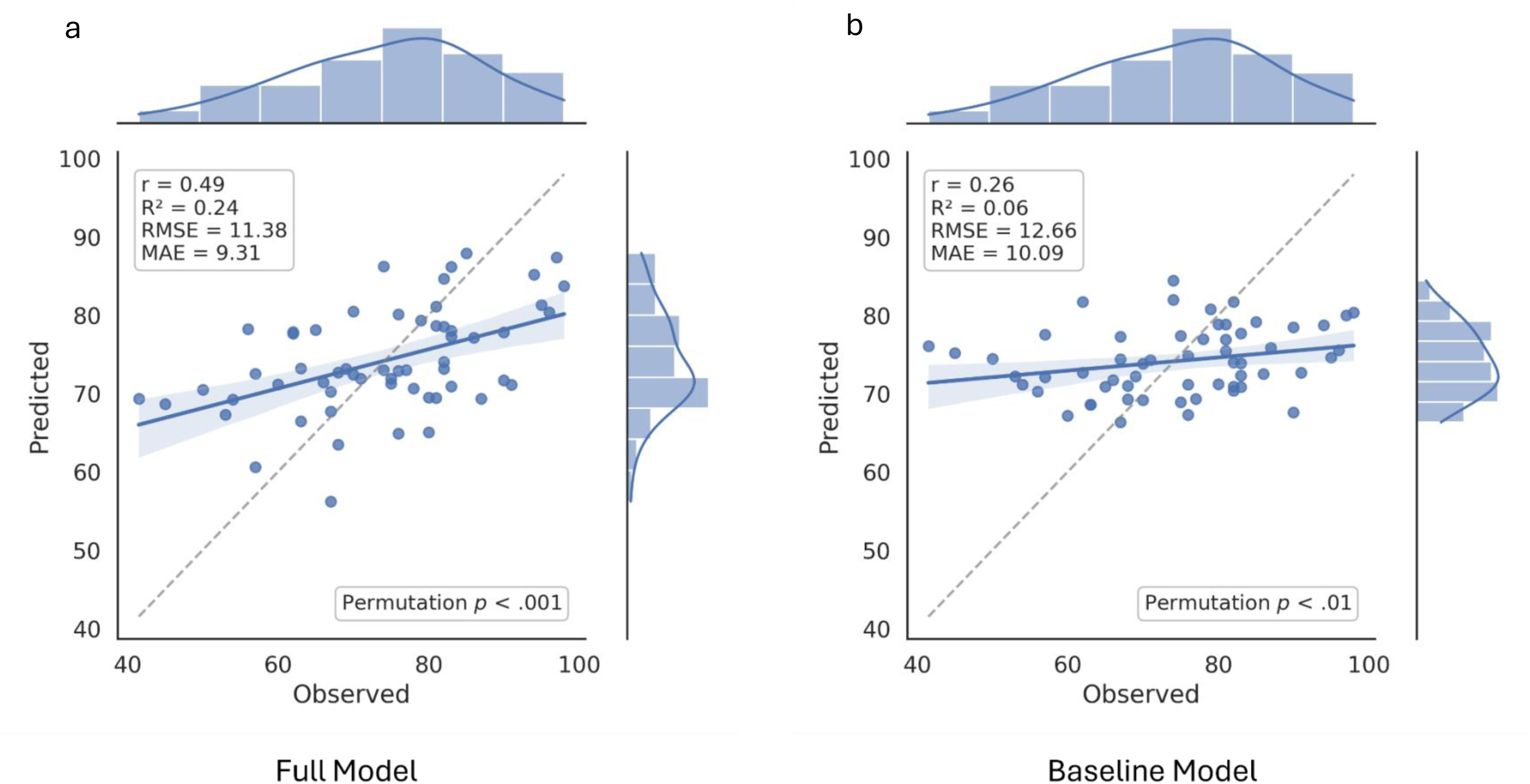
Relationship between observed and predicted BMRQ scores in older adults for the full model (a) and baseline model (b). *Note*. Full model (FA-MT-Sex model) = elastic-net model using FA features together with musical training score and sex as predictors; Baseline model = elastic net model using musical training score and sex as predictors. Each point represents one participant. The solid blue line indicates the least-squares regression fit between observed and predicted BMRQ scores, and the shade area represents the 95% confidence interval of the regression line. The dashed gray diagonal line represents perfect agreement between observed and predicted scores. Histograms and density curves along the top and right margins show the distribution of observed and predicted BMRQ scores, respectively. *r*, Pearson’s correlation coefficient; R², coefficient of determination; RMSE, root mean squared error; MAE, mean absolute error; Permutation *p*, *p* value of permutation tests.

**Figure S3a.**
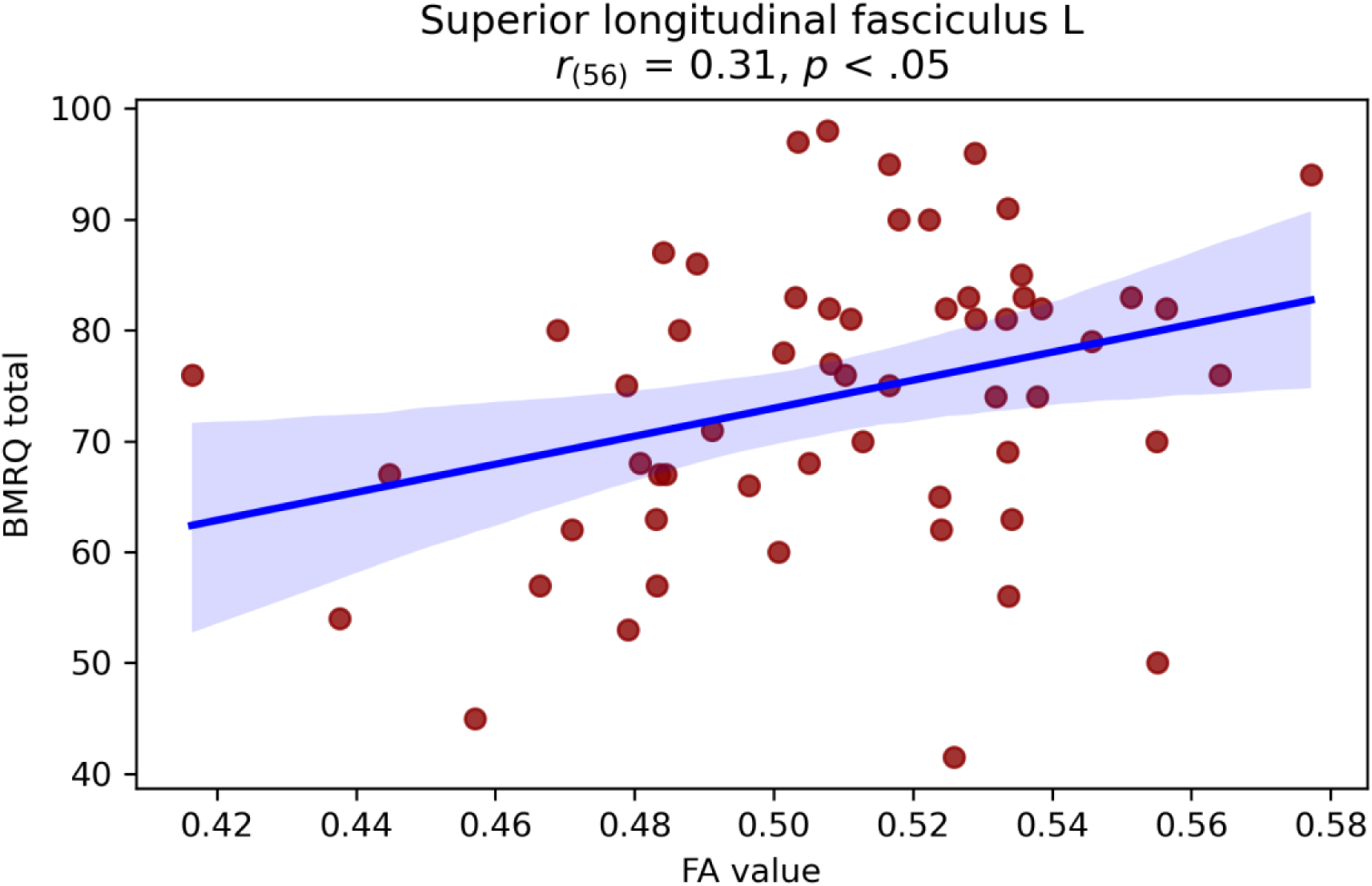
Correlation between FAs of the left superior longitudinal fasciculus tract (SLF) and BMRQ score. *Note.* Superior longitudinal fasciculus L, the left superior longitudinal fasciculus; BMRQ total, the total score of Barcelona Music Reward Questionnaire.

**Figure S3b.**
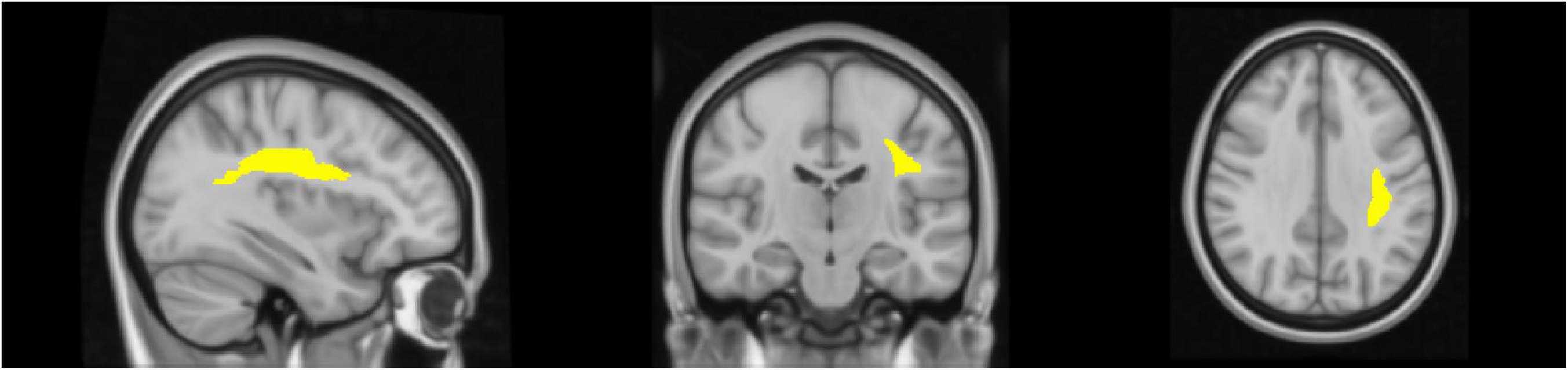
The illustration of the left superior longitudinal fasciculus (SLF) tract.

**Figure S4.**
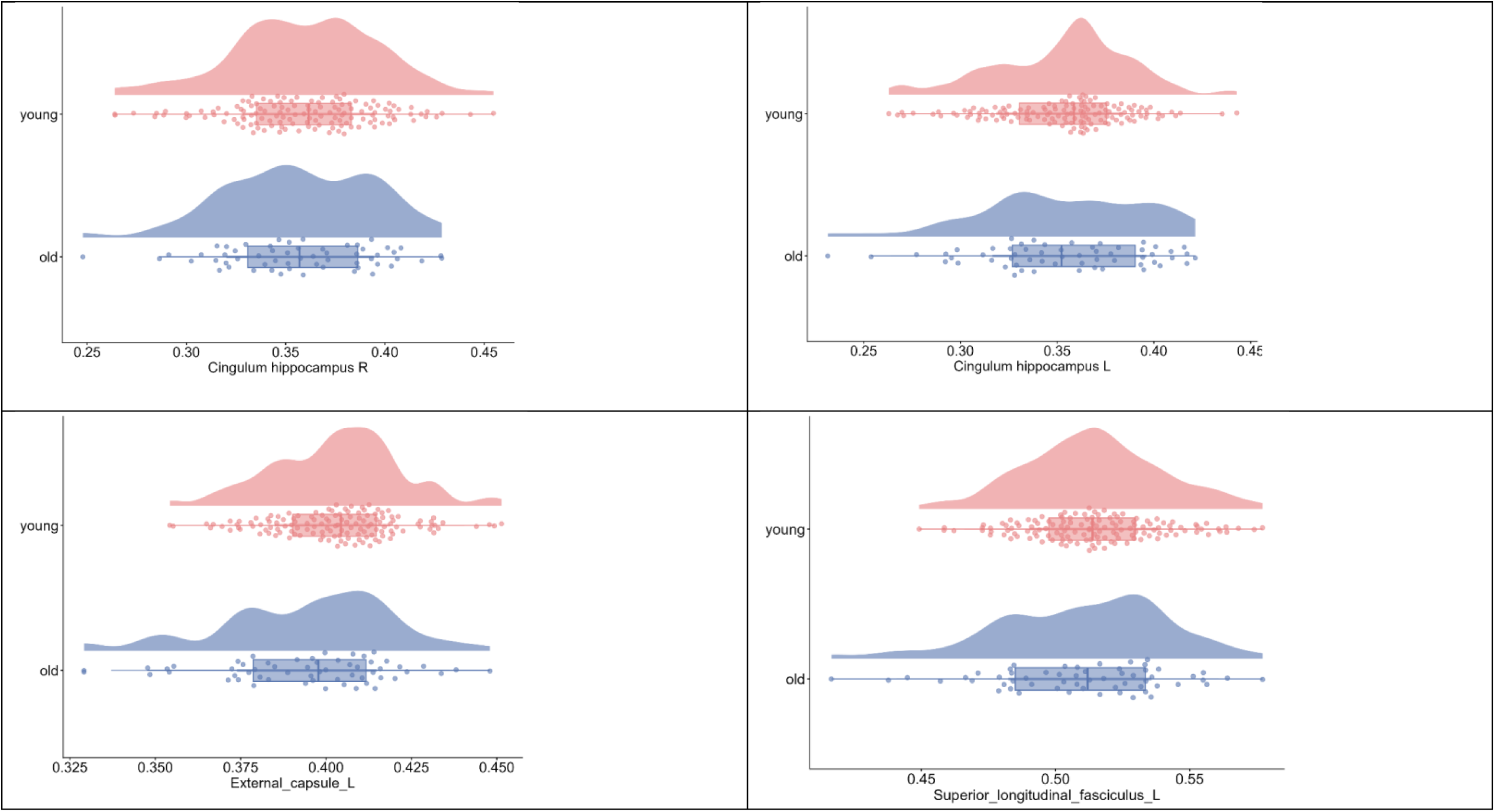
Distributions of FA values in the bilateral cingulum hippocampus, the left external capsule, and the left superior longitudinal fasciculus in older and young adults group.

